# Spatial distribution of extensively drug-resistant tuberculosis (XDR-TB) patients in KwaZulu-Natal, South Africa

**DOI:** 10.1101/132886

**Authors:** Thandi Kapwata, Natashia Morris, Neel R. Gandhi, Angela Campbell, Thuli Mthiyane, Primrose Mpangase, Kristin N. Nelson, Salim Allana, James C.M. Brust, Pravi Moodley, Koleka Mlisana, N. Sarita Shah

**Affiliations:** Environment and Health Research Unit, South African Medical Research Council, Johannesburg (South Africa); Biostatistics Unit, South African Medical Research Council, KwaZulu-Natal (South Africa); Emory University Rollins School of Public Health, Atlanta, GA (USA); University of KwaZulu-Natal, Durban, KwaZulu-Natal (South Africa); Albert Einstein College of Medicine and Montefiore Medical Center, Bronx, NY (USA); U.S. Centers for Disease Control and Prevention, Atlanta, GA (USA)

## Abstract

**Background:** KwaZulu-Natal province, South Africa, has among the highest burden of XDR-TB worldwide with the majority of cases occurring due to transmission. Poor access to health facilities can be a barrier to timely diagnosis and treatment of TB, which can contribute to ongoing transmission. We sought to determine the geographic distribution of XDR-TB patients and proximity to health facilities in KwaZulu-Natal.

**Methods:** We recruited adults and children with XDR-TB diagnosed in KwaZulu-Natal. We calculated distance and time from participants’ home to the closest hospital or clinic, as well as to the actual facility that diagnosed XDR-TB, using tools within ArcGIS Network analyst. Speed of travel was assigned to road classes based on Department of Transport regulations. Results were compared to guidelines for the provision of social facilities in South Africa: 5km to a clinic and 30km to a hospital.

**Results:** During 2011–2014, 1027 new XDR-TB cases were diagnosed throughout all 11 districts of KwaZulu-Natal, of whom 404 (39%) were enrolled and had geospatial data collected. Participants would have had to travel a mean distance of 2.9 km (CI 95%: 1.8-4.1) to the nearest clinic and 17.6 km (CI 95%: 11.4-23.8) to the nearest hospital. Actual distances that participants travelled to the health facility that diagnosed XDR-TB ranged from <10 km (n=143, 36%) to >50 km (n=109, 27%). The majority (77%) of participants travelled farther than the recommended distance to a clinic (5 km) and 39% travelled farther than the recommended distance to a hospital (30 km). Nearly half (46%) of participants were diagnosed at a health facility in eThekwini district, of whom, 36% resided outside the Durban metropolitan area.

**Conclusions:** XDR-TB cases are widely distributed throughout KwaZulu-Natal province with a denser focus in eThekwini district. Patients travelled long distances to the health facility where they were diagnosed with XDR-TB, suggesting a potential role for migration or transportation in the XDR-TB epidemic.

## BACKGROUND

Tuberculosis (TB) remains a major health burden globally [1], with 10.4 million cases estimated to have occurred in 2015 [2]. South Africa has the sixth highest burden of TB in the world (estimated 454,000 cases), and the second highest estimated incidence (834 per 100,000 population) [2]. Extensively drug resistant tuberculosis (XDR-TB) is defined as resistance to at least isoniazid, rifampicin, a fluoroquinolone and a second-line injectable drug, the most effective first- and second-line drugs for treating TB. XDR-TB has been reported in 105 countries and South Africa has the highest prevalence of XDR-TB in sub-Saharan Africa [2, 3]. Within South Africa, KwaZulu-Natal province has among the highest burden of XDR-TB, with 1,596 diagnosed cases in 2012 [4-6] and high rates of HIV co-infection (>90% among XDR-TB patients) [7].

Drug-resistant TB is characterized by delayed diagnosis, in part due to policies that restrict drug-susceptibility testing. Treatment success rates are low. Consequently, patients experience prolonged infectious periods, which increases the risk of ongoing transmission to others. Delays in diagnosis also place patients at risk for clinical decline and increased mortality rates, particularly in the setting of HIV co-infection [7, 8]. Furthermore, adults suffering from TB lose several months of working time, resulting in a decrease in household income [9], further underscoring the need for early diagnosis and initiation of effective treatment. However, both are dependent on the access that individuals have to health facilities offering appropriate TB diagnosis and treatment services.

Access to health services is difficult to define because it encompasses several factors, including quality of care or service, geographical accessibility, suitability of available services and financial constraints of patients and health systems. Of these, geographic accessibility is a crucial barrier to individuals accessing health facilities and is important to understand as programs expand TB services. Several studies in developing countries have provided evidence that physical proximity of healthcare services can play an important role in the use of healthcare facilities [10-12]. Recent advances in geographical information systems (GIS) software that facilitate distance-based measures focused on distance and travel time have led to more accurate analysis of driving distance and travel time based on actual road networks[13, 14].

This study aimed to describe the geographic distribution of patients with XDR-TB in relation to health facilities. Specifically, we sought to assess ease of access based on travel times and distances to the closest health facility, as well as to the actual health facility at which XDR-TB was diagnosed. This analysis provides an understanding of the availability and utilisation of public healthcare services among XDR-TB patients throughout the province.

## MATERIALS AND METHODS

### Study area

KwaZulu-Natal province comprises 11 districts (Figure 1) and has a population of 10.3 million persons, 53% of whom reside in rural areas[15]. KwaZulu-Natal harbours nearly half of South Africa’s XDR-TB burden and has more than 1.6 million people living with HIV. The province has 56 hospitals and 562 clinics [16] whose distribution mirrors the overall population distribution, with 57% located in rural areas[15].

**Figure 1:**
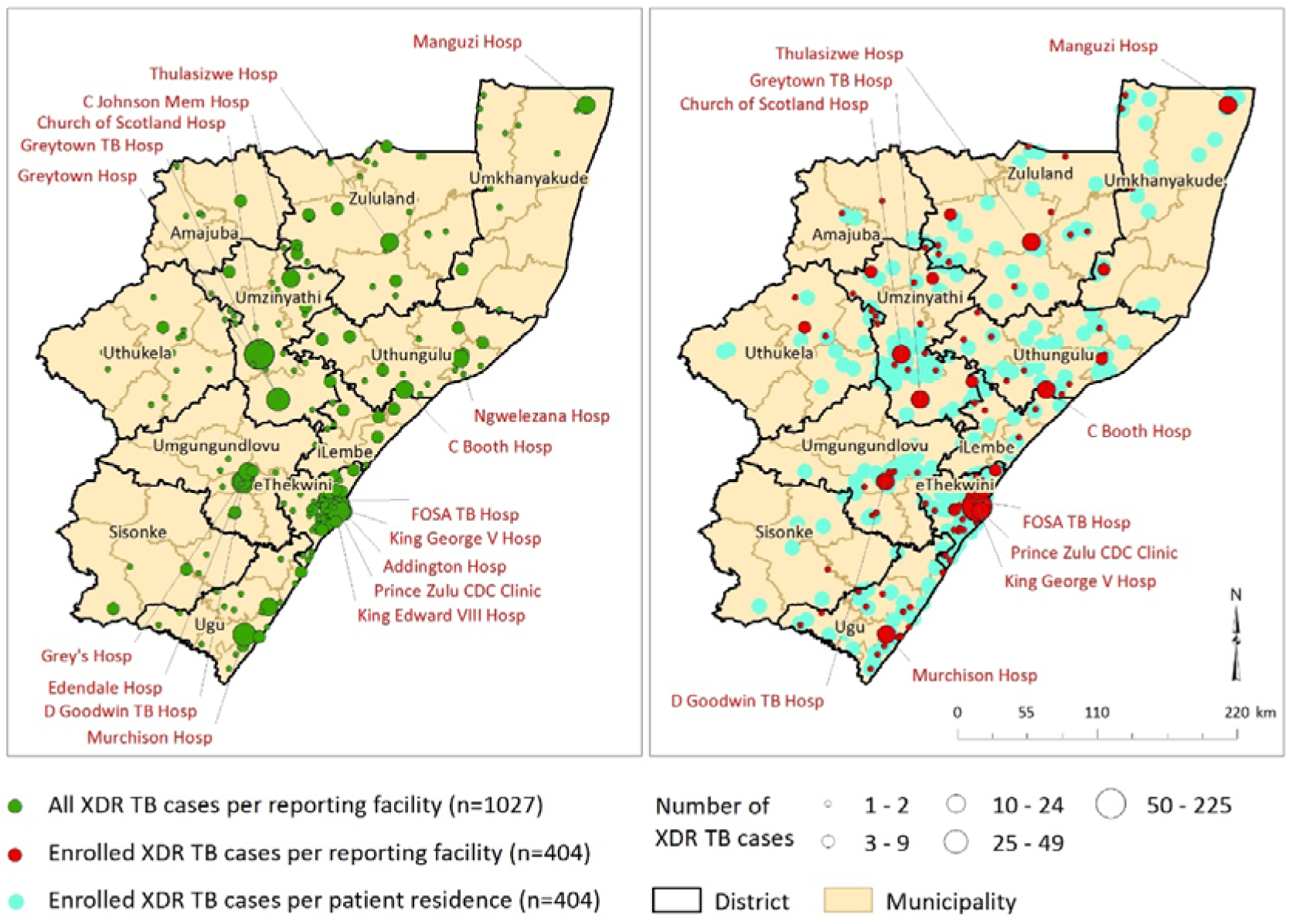
Distribution of all XDR-TB cases diagnosed per health facility (left panel) and enrolled XDR-TB cases per health facility and residential location (right panel), KwaZulu-Natal, 2011–2014.

### Study subjects and data collection

The study prospectively enrolled culture-confirmed XDR-TB patients diagnosed from May 2011 through August 2014, residing in KwaZulu-Natal. A single referral laboratory conducts drug-susceptibility testing (DST) for all public healthcare facilities in the province, allowing for complete capture of all diagnosed XDR-TB cases. For deceased or severely ill participants, consent was obtained from next-of-kin. Interviewers collected information about socio-demographics, TB and HIV history, and location and duration of all hospitalisations in the preceding 5 years. A global position system (GPS) coordinate location was collected at each participant’s home and the diagnosing facility was identified as the clinic or hospital from which the specimen that grew XDR-TB was submitted to the provincial lab.

### Analysis of distance to closest healthcare facility and diagnosing facility

ArcGIS Network Analyst is an extension within ArcGIS® software [17] that allows users to model realistic road network conditions by incorporating road network data, lengths of road segments, speed limits, turn restrictions and vehicle height restrictions. Network Analyst uses Dijkstra’s Algorithm to solve routing problems, which is based on distance and time criteria [18]. The assumption made in this analysis is that travel to specified locations is along the shortest route in the network. Estimated travel times and distances do not account for delays which may be encountered along the road network for various reasons.

Travel distances of study participants were determined to the nearest clinic and hospital. Road network data including road classes were utilised for this analysis [19]. The “closest facility” tool was used to obtain distance and the “origin-destination (OD) matrix” tool was used to obtain time, both tools available within the ArcGIS® Network Analyst extension. The closest facility tool measures the time of traveling between incidents (location of study participants) and facilities (health facilities) and determines which are nearest to one other. The OD cost matrix finds the least-cost path (measured by time) from each origin to the nearest specified destination. Different speeds were assigned to various road classes within the network; speed assigned per segment was determined by the KwaZulu-Natal Department of Transport regulations [20]. Table 1 shows the hierarchy of road classes and their associated speeds.

**Table 1:** Road classes and road speeds in KwaZulu-Natal province, as defined by the KwaZulu-Natal Department of Transport.

| Road classification | Movement network | Speed<br>(km/h) |
| --- | --- | --- |
| Regional distributor | Vehicle-only route | 120 |
| Primary distributor | Vehicle-only route | 100 |
| District distributor | Mixed pedestrian and vehicle route | 80 |
| Local distributor | Mixed pedestrian and vehicle route | 60 |
| Access street | Mixed pedestrian and vehicle route | 40 |

The quality of road network data is crucial to the analysis providing accurate results, therefore care was taken to ensure that all polylines comprising the network did not overlap, were not duplicated, and did not have breaks in connectivity (end points of lines that are not joined to other lines in the network). Data cleaning and quality checking was conducted using a suite of geoprocessing tools within ArcGIS®.

Residential location of the participant and location of the health facility that sent the diagnostic XDR-TB sputum were both known for participants enrolled into the study, and were used to determine actual travel distance and time per participant. The locations of health facilities to which study participants reported were considered in relation to the cleaned, coded provincial road network in KwaZulu-Natal, with road features divided into four main classes: regional, primary, district and local [20]. The travel distance and time from the participant’s residence to the actual health facility at which he/she presented was calculated using the ArcGIS® “XY to Line” tool.

### Ethical Considerations

The study was approved by the Institutional Review Boards of Emory University, Albert Einstein College of Medicine, and the University of KwaZulu-Natal, and by CDC’s National Center for HIV, Hepatitis, STDs and Tuberculosis.

## RESULTS

A total of 1,027 XDR-TB cases were diagnosed at health facilities in KwaZulu-Natal between 2011 and 2014, of which 521 (51%) were screened and 404 (38%) were enrolled into the study. Enrolled cases were distributed across all eleven districts in the province (Figure 1).

Participants would have had to travel a mean distance of 2.9 km (CI 95%: 1.8-4.1) to the nearest clinic and 17.6 km (CI 95%: 11.4-23.8) to the nearest hospital (Table 2). Participants enrolled from the predominantly rural districts of uThukela (69.7% rural; 38.9km), uMkhanyakude (94.4% rural; 20.2km), Ugu (82.4% rural; 23.1km) and Sisonke (79.7% rural; 24.6km) would have had to travel the farthest distance to their nearest hospital, all greater than the overall provincial mean. The guidelines provided by the Council for Scientific and Industrial Research (CSIR) for the provision of social facilities in South Africa sets the acceptable travel distance at 5 km to a clinic and 30 km to a hospital[21]. Two districts exceeded this recommendation, with participants in Sisonke residing an average travel distance of 6.4 km from the nearest clinic and participants in uThukela residing 38.9 km away from the nearest hospital (Table 2).

**Table 2:** Mean travel distance and time to the nearest clinic, nearest hospital and the facility that diagnosed XDR-TB, by district.

| District | n | Travel distance to nearest facility (kms) |  | Travel time to nearest facility (hrs) |  | Proportion Population |  | Travel distance to actual facility (kms) | Travel time to actual facility (hours) |
| --- | --- | --- | --- | --- | --- | --- | --- | --- | --- |
|  |  | Clinic | Hospital | Clinic | Hospital | Urban | Rural |  |  |
| Sisonke | 6 | 6.37 | 24.55 | 0.08 | 1.02 | 20.3 | 79.7 | 119.93 | 1.50 |
| Zululand | 31 | 4.89 | 17.36 | 0.12 | 0.80 | 19.5 | 80.5 | 72.12 | 0.90 |
| uMkhanyakude | 22 | 3.63 | 20.24 | 0.12 | 2.86 | 5.6 | 94.4 | 89.91 | 1.12 |
| uMzinyathi | 65 | 3.63 | 14.42 | 0.11 | 1.16 | 17.4 | 82.6 | 42.99 | 0.54 |
| Uthungulu | 36 | 2.82 | 9.5 | 0.05 | 2.05 | 18.1 | 81.9 | 201.37 | 2.52 |
| uThukela | 17 | 2.73 | 38.87 | 0.08 | 0.85 | 30.3 | 69.7 | 50.73 | 0.63 |
| Ugu | 37 | 2.47 | 23.06 | 0.05 | 1.43 | 17.6 | 82.4 | 174.54 | 2.18 |
| uMgungundlovu | 39 | 2.23 | 18.9 | 0.06 | 0.25 | 58.1 | 41.9 | 36.11 | 0.45 |
| iLembe | 15 | 2.09 | 9.32 | 0.04 | 1.81 | 36.5 | 63.5 | 45.59 | 0.57 |
| Amajuba | 5 | 0.74 | 11.6 | 0.27 | 0.95 | 55.3 | 44.7 | 201.39 | 2.52 |
| eThekwini | 131 | 0.49 | 6 | 0.01 | 0.73 | 84.8 | 15.2 | 19.57 | 0.24 |
| TOTAL | 403 | 2.92 | 13.92 | 0.06 | 1.11 | 33 | 67 | 68.82 | 0.86 |

Mean distances that study participants actually travelled to the diagnosing health facility are described for each of the 11 districts and 47 municipalities in KwaZulu-Natal in Figure 2. Study participants residing in Amajuba, Uthungulu, Sisonke and Ugu distrits travelled the farthest, on average, to the facility at which they were diagnosed (>100 km). Participants in Ukhanyakude, Zululand and Uthukela also travelled large distances (50-100km), while those in the central districts (Umzinyathi, Umgungundlovu and eThekwini) travelled the lowest mean distances (>10-50km). The highest proportion of patients seeking care outside their district of residence (>60%) came from Sisonke, iLembe and Amajuba districts (83%, 80% and 67%, respectively).

**Figure 2:**
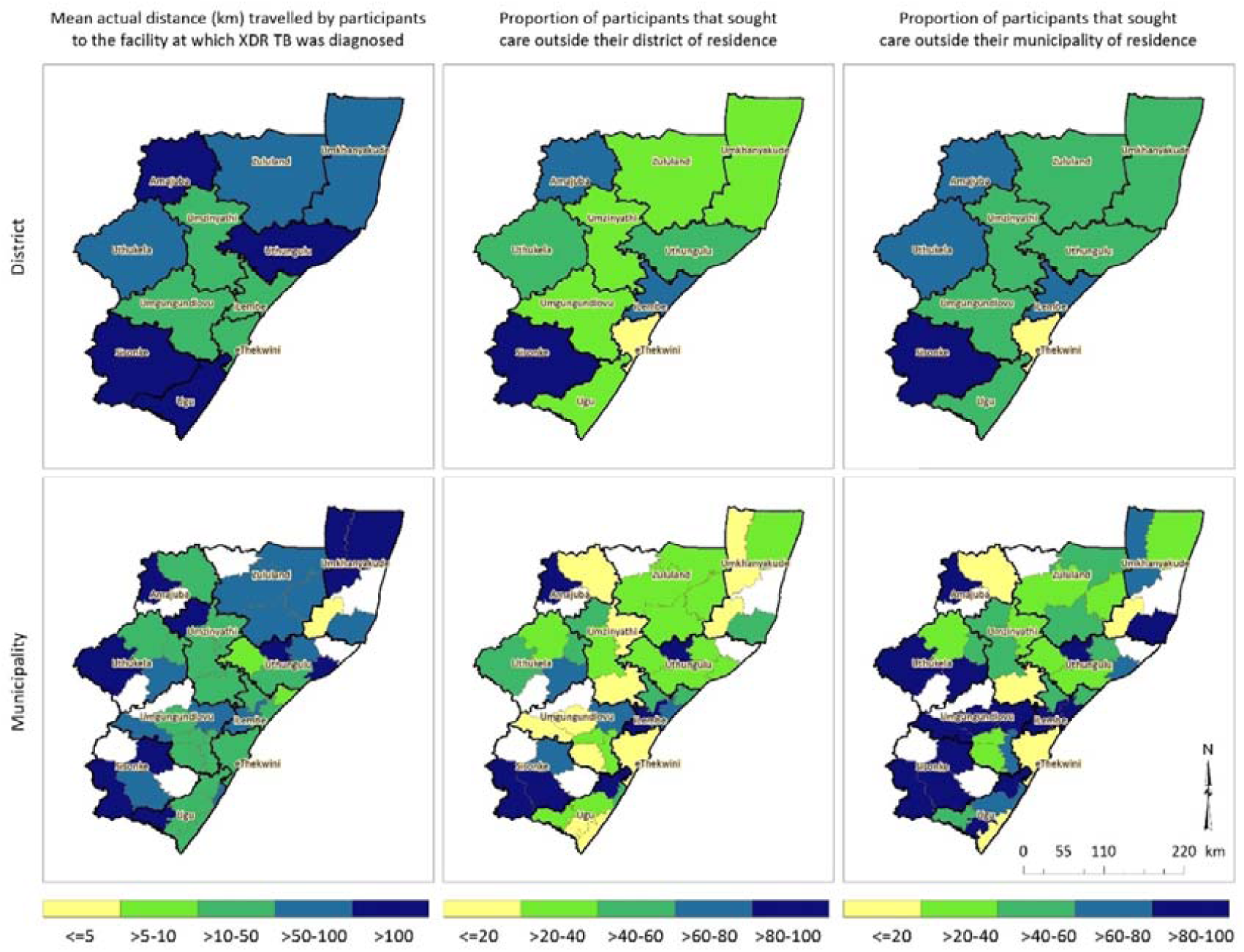
Mean actual distance travelled by participants to the health facility at which XDR-TB was diagnosed, mean distance travelled farther than the nearest clinic, and proportion that sought care outside their district and municipality of residence, districts and municipalities of KwaZulu-Natal, 2011–2014.

Of the 404 participants, 111 (28%) reported to facilities outside their district of residence, and 185 (46%) reported to a health facility in the eThekwini District; 36% of these (16% of all enrolled cases) originated outside of the Durban metropolitan area, but were diagnosed with XDR-TB in Durban. Actual distances that participants travelled to the health facility at which they were diagnosed ranged from < 10 km (n=143, 36%) to > 50 km (n=109, 27%) (Table 3).

**Table 3:**
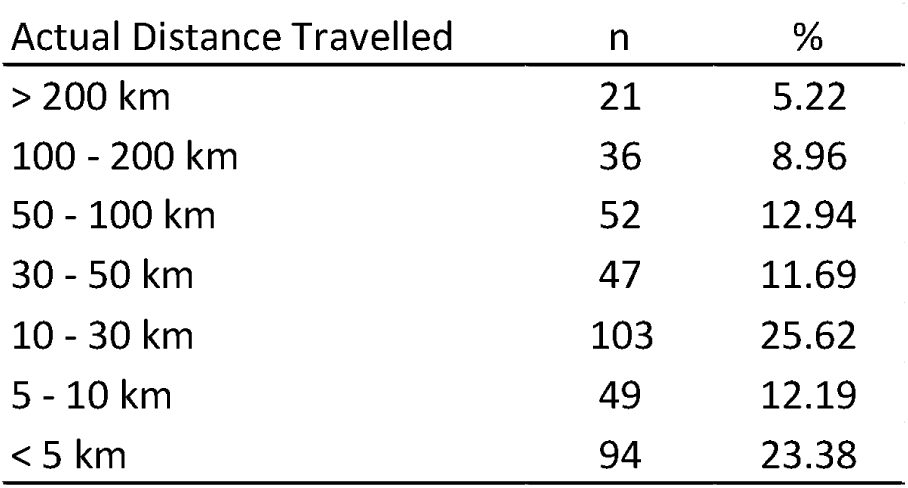
Actual distance travelled by participants (n=404) to the health facility at which XDR-TB was diagnosed, KwaZulu-Natal, 2011–2014.

| Actual Distance Travelled | n | % |
| --- | --- | --- |
| > 200 km | 21 | 5.22 |
| 100 - 200 km | 36 | 8.96 |
| 50 - 100 km | 52 | 12.94 |
| 30 - 50 km | 47 | 11.69 |
| 10 - 30 km | 103 | 25.62 |
| 5 - 10 km | 49 | 12.19 |
| < 5 km | 94 | 23.38 |

The average travel distances significantly exceed the distances that individuals would have had to travel to the facility nearest to their homes. A spider plot illustrating straight-line distances between participant residence and the facility at which they were diagnosed further confirmed that study participants sought care at health facilities distant from their place of residence (Figure 3).The majority (77%) of participants travelled farther than the recommended distance to a clinic (5 km) and 39 % travelled farther than the recommended distance to a hospital (30 km).

**Figure 3:**
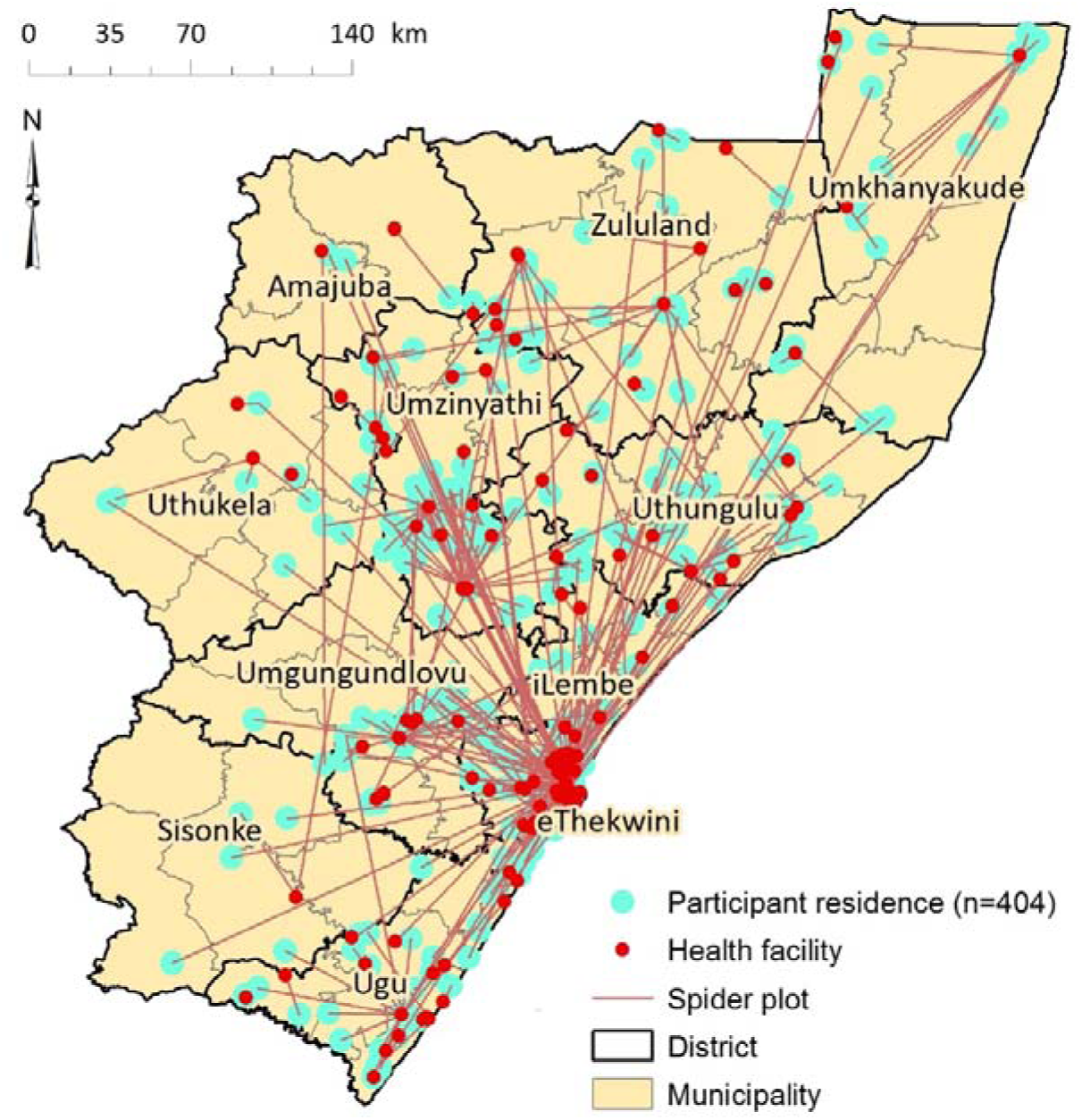
Spider plot showing straight line distance from participants’ residence to the health facility at which they presented that diagnosed XDR-TB, KwaZulu-Natal, 2011–2014.

## DISCUSSION

South Africa is experiencing a widespread epidemic of XDR-TB, and has among the highest rates of XDR-TB worldwide. We have previously shown that the majority of XDR-TB cases occur due to person-to-person transmission of XDR-TB strains [22]. Prolonged infectious periods in XDR-TB patients may be caused by delayed diagnosis, delayed treatment, or both, which leads to ongoing transmission. In the current study, we assessed the distance to the diagnosing health facility in a cohort of patients with XDR-TB and found that a large proportion had sought health care outside their district of residence, with 27% traveling over 50km to a facility for XDR-TB diagnosis. Almost half (46%) of participants were diagnosed at a health facility in eThekwini district, of whom 36% resided outside the Durban metropolitan area. Together, these findings suggest an important role for within-province travel for healthcare in the ongoing XDR-TB epidemic.

A major strength of this study is that it enrolled participants from all 11 districts in KwaZulu-Natal province and used the most up-to-date road network data to model travel times and distances. Furthermore, road network analysis was conducted using GPS coordinate location of the participant’s residential address in relation to both the physical location of health facilities in closest proximity, and the physical location of the actual facility that sent the diagnostic XDR-TB sputum. Similar studies have attempted to model accessibility of health care facilities using road networks in South Africa. Tanser, Gijsbertsen [23] collected data on methods of accessing health care from 23,000 homesteads in Hlabisa municipality within uMkhanyakude district. They found that the average travel times using public transport were 81 and 170 minutes to a clinic and hospital, respectively. However, the study did not calculate travel *distance* and it was limited to one district in KwaZulu-Natal province. Nteta, Mokgatle-Nthabu [24] did not undertake road network analysis, but surveyed participants who attended three community health centres (CHCs) in Gauteng province. Their findings revealed that a large proportion of respondents (71%) travelled 30 minutes or less to the nearest clinic and a smaller proportion (24%) travelled distances between 30 minutes and an hour. The results of their distance analysis found 45% of study participants travelled less than 5km and 39% travelled 5 to 10km. However, that study did not model travel based on road networks, the sample size was smaller (235 participants) and it was confined to a single district (Tshwane).

We found that study participants would have had to travel 2.9 km and 17.6 km, on average, from their residence to the nearest public health clinic and hospital, respectively. However, they travelled an average of 96 km to the actual facility (both clinics and hospitals combined) that diagnosed XDR-TB disease. This distance significantly exceeds the average distance reported in Tshwane district of between 5 and 10 km [24]. The mean time to the health facility was found to be 1.2 hours, considerably greater than the mean time of 30.7 minutes reported by Harris, Goudge [25]] who aimed to assess inequalities in access to healthcare in South Africa. Districts with large proportions of their population living in rural areas had longer travel distances for participants, both to their nearest facility and to the facility at which they actually presented. Access to public transportation and poor quality of road networks may contribute to increased travel distance and time to health facilities in rural areas.

Study participants travelled substantially farther than their nearest facilities to seek care. Reasons for this health seeking behaviour are likely to vary, and may include patients seeking care in areas where they are employed instead of where they reside; patients seeking care at hospitals in urban areas that are perceived to have favourable reputations in communities; patients wanting to seek care at facilities that provide better quality of care in terms of available treatments, facilities, medication and staff; as well as patients wanting to use health care facilities that have a shorter waiting time and queues [25]. Other factors that could be contributing to patients travelling long distances for care could be the stigma associated with TB in South Africa[26-28]. Whatever the reasons for seeking care at facilities farther away, time losses resulting from travelling long distances are a potential threat to TB diagnosis, treatment initiation and retention, all of which contribute to ongoing transmission [29].

Long travel distances to access public health care has additional consequences of greatly increasing the possibility of transmission of XDR-TB at the community level. Among study participants who reported regularly traveling long distances in the months preceding their XDR-TB diagnosis, 80% (44 of 55) reported using public transportation when traveling; these modes of transportation included combis, taxis or buses that are crowded, poorly-ventilated vehicles. In addition, although not directly measured, patients are likely to have sought care at their local health facility before traveling to a farther facility for XDR-TB diagnosis and care. Taken together, each patient with XDR-TB may be spending a minimum of 2-4 hours traveling to and from health facilities while infectious (i.e., before a diagnosis is made and effective treatment is started), creating numerous opportunities for transmission *en route*.

Our results showed that 43% of the enrolled participants sought health care at eight specialised TB hospitals that are provincial facilities; the remainder were distributed between standard hospitals and clinics, mobile clinics and community health centres. These specialised hospitals serve communities with an elevated incidence of TB and they provide long-term inpatient care for patients with chronic TB. However, the fact that only eight such facilities provincially reported the plurality of the XDR-TB cases highlights the shortage of these specialised facilities, perhaps a factor compelling patients to travel long distances. New rapid molecular diagnostics for drug resistance that are situated closer to patients, combined with decentralized models of care may reduce delays in diagnosis and treatment, and reduce transmission [30, 31].

A limitation of this study was that all participants were assumed to have access to either public or private modes of transport, which might not have been the case. We also calculated uninterrupted travel along the roads to obtain travelled distances and time. Therefore, we did not account for any factors that would affect flow of traffic or slow vehicles down. Distances were calculated from a single home address, which may not represent the starting point for travel to all health facilities in this population. Furthermore, we did not obtain qualitative information from participants about their reasons for seeking health care far from their residence; nor did we compare the capacity for providing adequate care at the closest clinic or hospital to the capacity at the actual diagnosing facility. However, the primary capacity that would be needed is clinical suspicion for TB, and sputum sample collection and transport for drug resistance testing at a referral laboratory.

Transmission of XDR-TB is driving the epidemic in South Africa. Our findings demonstrate the long distances a large proportion of patients travelled for care that correctly diagnosed XDR-TB. The role of migration has been well-studied and documented for the HIV epidemic in South Africa, but far less is known about the impact of migration on TB and XDR-TB transmission. As people move for work, school, family or health reasons throughout KwaZulu-Natal, a better understanding of where transmission is occurring can help guide interventions aimed at halting ongoing XDR-TB spread.

## ACKNOWLEDGMENTS

### Acknowledgments

We are grateful to the study teams at the University of KwaZulu-Natal and South Africa Medical Research Council for their tireless efforts in data collection, record abstraction, participant recruitment and interviews. We thank the participants and their families who consented to participate in this study.

### Funding Source

This study was primarily funded by a grant from the US National Institute of Allergy and Infectious Diseases (NIAID), National Institutes of Health (NIH): R01AI089349 (PI Gandhi). It was also supported in part by NIH/NIAID grants: R01AI087465 (PI Gandhi), K23AI083088 (PI Brust), K24AI114444 (PI Gandhi), Emory CFAR P30AI050409 (PI Curran), Einstein CFAR P30AI051519 (PI Goldstein), by Einstein/Montefiore ICTR UL1 TR001073 (PI Shamoon) and by Atlanta CTSI UL1 TR000454 (PI Stephens).

### Conflicts of Interest

No conflict of interest reported for any authors.

### Disclaimer

The findings and conclusions in this manuscript are those of the authors and do not necessarily represent the official position of the Centers for Disease Control and Prevention or the U.S. Department of Health and Human Services.

